# Widespread transcription initiation within coding sequences marks tissue identity and accessible chromatin

**DOI:** 10.1101/2024.03.27.587116

**Authors:** Xutong Wang, Jingbo Duan, Chancelor B. Clark, Wanjie Feng, Jianxin Ma

**Affiliations:** Department of Agronomy, Purdue University, West Lafayette, IN 47907, USA; Center for Plant Biology, Purdue University, West Lafayette, IN 47907, USA; National Key Laboratory of Crop Genetic Improvement, College of Plant Science and Technology, Hubei Hongshan Laboratory, Huazhong Agricultural University, Wuhan, Hubei 430070, China

## Abstract

Alternative transcription initiation (ATI) appears to be a ubiquitous regulatory mechanism of gene expression in eukaryotes, but the extent to which it affects the products of gene expression, and how it evolves and is regulated remain unknown. We first identified transcription start sites (TSSs) in eight soybean tissues using recently developed STRIPE-seq and then analyzed ATI in the context of tissue identity and chromatin architecture. We defined 193,579 TSS clusters/regions (TSRs) in 37,911 annotated genes, with 56.5% located in putative regulatory regions upstream of start codons and 43.5% from start codons to 3’ untranslated regions, which, together, are responsible for changes in open reading frames of 24,131 genes. Overall, duplicated genes possess more TSRs, exhibited lower degrees of tissue-specificity, and have undergone stronger purifying selection than singletons. Strikingly, 6,845 genes possess ATI within coding sequences (CDSs). These CDS-TSRs are highly tissue-specific, shorter than those located in canonical regulatory regions, and do not have TATA-boxes typical for the core promoters. Furthermore, the CDS-TSRs are embedded in nucleosome-free regions and flanked by nucleosomes with enhanced levels of active histone marks associated with transcriptionally active chromatin, suggesting that non-canonical ATI is epigenetically regulated and largely responsible for tissue-specific functions and tissue identity. Our study highlights the genomic and epigenomic factors shaping the distribution patterns and tissue-specificity of ATI in regulatory and coding sequences, as well as the significance of ATI in the alternation of proteins encoded by tissue-specifically expressed genes in the context of genome duplication and fractionation.

## Introduction

Transcriptional regulation is crucial for various biological processes such as development, tissue differentiation, and environmental responses (Singh 1998; Macrae and Long 2012). Understanding of the fundamental aspects of transcription relies on the accurate identification of transcription start sites (TSSs), which are typically hallmarked with cis-regulatory DNA elements, TATA-boxes, in various eukaryotic genomes (Davuluri et al. 2008; Mejía-Guerra et al. 2015; Policastro et al. 2020; Thieffry et al. 2020). The TSSs are core part of promoters and often clustered as Transcription Start Regions (TSRs) (Policastro et al. 2020). Canonical TSRs are typically located in the 5’ upstream regulatory regions of genes, but non-canonical TSRs besides those regions were also observed. Despite the availability of genome sequences and annotations for numerous eukaryotic organisms; however, TSRs in most of these organisms remain to be identified towards an in-depth and comprehensive understanding of the regulatory mechanisms of transcription and the functional consequences.

Over the past few decades, several methods, including cap analysis of gene expression (CAGE) (Murata et al. 2014), nano-CAGE (Batut and Gingeras 2013), paired-end analysis of TSSs (PEAT) (Shahmuradov et al. 2017), and RNA annotation and mapping of promoters for the analysis of gene expression (RAMPAGE) (Ni et al. 2010; Batut and Gingeras 2013; Napoli 2017) have been developed and extensively used to identify TSRs in eukaryotes. Each of these methods has its own advantages and disadvantages, but the major limitation is the costs and laboriousness (Shahmuradov et al. 2017; de Medeiros Oliveira et al. 2021). Recently, a fast, efficient, and simple protocol, called survey of transcription initiation at promoter elements with high-throughput sequencing (STRIPE-seq), has been developed and first used for genome-wide identification of the capped 5’ ends of transcripts in yeast and human (Policastro et al. 2020). STRIPE-seq enables the identification of TSRs and quantification of gene expression levels simultaneously, representing an additional advantage compared with other methods.

Soybean (*Glycine max* [L.] Merr.) is a primary source of protein for livestock feed as well as a valuable contributor to vegetable oil and various industrial and bioenergy compounds (Hartman et al. 2011). As such, soybean is among the few crops whose reference genomes were assembled over a decade ago (Schmutz et al. 2010). One of the unique features of the soybean genome is that it has experienced two recent rounds of whole-genome duplication (WGD) events that occurred ∼59 and ∼5-13 million years ago (MYA), respectively, followed by the loss of a large proportion of the duplicated genes (Schmutz et al. 2010; Du et al. 2012; Zhao et al. 2017). On the other hand, although approximately two-thirds of the duplicated genes formed from the more recent WGD event are retained in the current soybean genome, the majority of the duplicated genes showed distinct expression patterns in terms of relative abundance of transcripts and tissue specificity, which possibly reflect their sub-functionalization and/or neo-functionalization (Zhao et al. 2017). Therefore, soybean is an ideal system to understand how the transcription patterns such as the distribution of TSSs of duplicated genes have been shaped by WGD and subsequent genomic and epigenomic differentiation.

The annotation of putative TSSs in the soybean reference genome was largely done based on prediction without experimental validation. In this study, STRIPE-seq was used to identify and validate TSRs in the soybean reference genome, heralding the inaugural application of this technology within plant systems. Through profiling TSRs in the entire genome across five vegetative tissues and three reproductive tissues, we were able to identify alternative transcription initiations (ATIs) in the context of tissue specification, subgenome fractionation, and epigenomic features, revealing the evolutionary factors and their interplay driving transcriptional and functional innovation of plant genes in a paleopolyploid genome.

## Results

### STRIPE-seq identified genome-wide TSRs in soybean

We identified TSRs in eight soybean tissues including leaves, stems, stem tips, roots, nodules, flowers, pods, and developing seeds, using the STRIPE-seq protocol, with slight modification involving the incorporation of the RiboMinus Plant Kit (Invitrogen) for rRNA depletion from total RNAs isolated from each of the eight tissues (Policastro et al. 2020). This modification was important as the terminator exonuclease (TEX) included in STRIPE is less effective than “RiboMinus” at removing rhizobial rRNAs from the nodule tissue that contains procaryotic cells (Čuklina et al. 2016). The STRIPE-seq of the eight issues generated 646 million reads (**Supplemental Table S1**). On average, 92% of the reads from each tissue contained a unique molecular identifier (UMI), enabling us to filter out redundant reads created by PCR (**Supplemental Table S1**). Only non-redundant reads uniquely mapped to the soybean genome were included for subsequent analyses (**Fig. 1a**). Saturation analysis indicated that the read counts for all eight tissues were sufficient for TSR identification (**Supplemental Fig. S1**). Our data revealed that the median TSRs were centered at 42 bp downstream of the TSSs annotated in the reference genome (**Fig. 1b and Supplemental Fig. S1**).

**Fig. 1.**
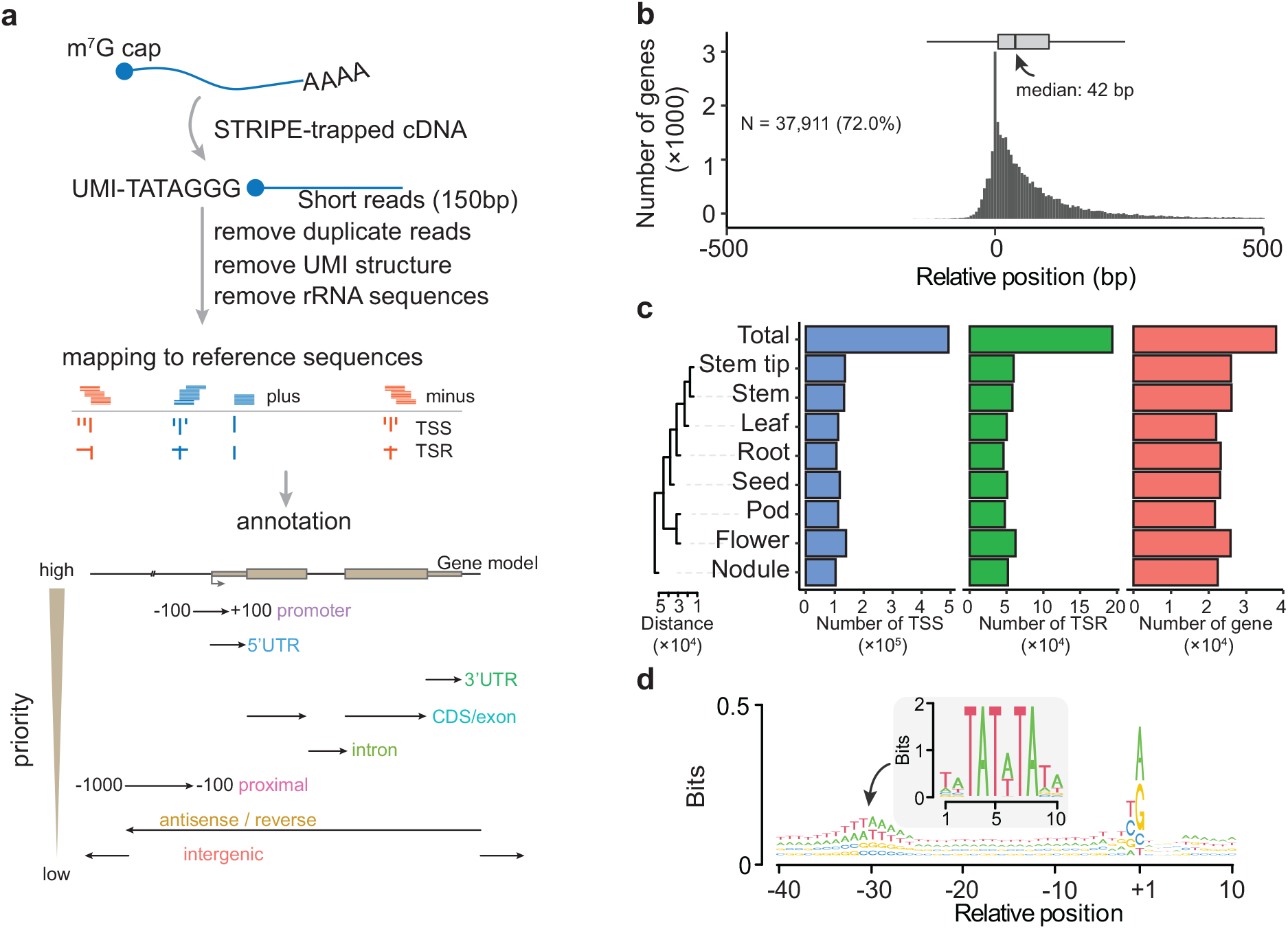
Annotation and evaluation of TSRs detected by STRIPE-seq. **a,** Workflow of processing STRIPE-seq reads and hierarchical annotation strategy. Schematic representation of the annotation of TSRs to the reference genome. The priority annotation levels from high to low are shown on the left side of the bottom panel. **b,** The distribution of TSRs compared with annotated TSS (Williams 82 version 2). The boxplot at the top represents the distribution of distances of TSRs between annotated TSSs and the detected TSRs. **c,** Statistics of the number of TSSs, TSRs, and covered genes in eight tissues and total. The left side cladogram represents the distance among tissues with the Euclidean metric using the relative abundance of TSRs. **d,** Sequence patterns around all unique TSSs. The TSS was defined as the peak site of the TSR. The motif in the central panel represents the TATA-box calculated in soybean.

A total of 492,858 unique TSSs, which defined 193,579 TSRs of 37,911 (72%) genes annotated in the reference genome, were defined (**Fig. 1c**). TSR numbers vary considerably among the eight tissues, ranging from 45,943 in roots up to 62,214 in flowers, with an average of 53,354 per tissue. Likewise, the average numbers of genes with defined TSRs range from 21,998 in leaves up to 25,806 in flowers, with an average of 23,760 per tissue, and the nodule tissue exhibited the most unique TSRs compared with other tissues (**Fig. 1c**). To evaluate the accuracy of TSRs defined by STRIPE-seq, we compared all the TSSs identified in this study with those annotated in the reference genome, those suggested by *de novo* RNAseq transcripts, and those defined by full-length RNAseq. The TSRs defined by STRIPE-seq uniquely identified both TATA-box and Pyrimidine-Purine (PyPu) motifs in the Initiator (*Inr*) element (**Supplemental Fig. S2**), which were enriched at at +30 and +1 bp upstream of the identified TSRs (**Fig. 1d**). Our results underscore the efficacy of STRIPE-seq for identifying TSRs in soybean.

### Genes with multiple TSRs experienced stronger purifying selection than those with a single TSR

The identified 193,579 TSRs varied drastically in length, ranging from a single nucleotide to hundreds of base pairs (**Fig. 2a and Supplemental Table S2**) and were categorized into three types based on their shapes defined by interquartile range (IQR) (**Fig. 2b**): single-nucleotide TSRs (S), narrow TSRs (N, IQR ≤ 4), and broad TSRs (B, IQR > 4) (Thodberg et al. 2019), accounting for 64.8%, 16.3%, and 18.9%, respectively (**Fig. 2a**). Overall, the genes with “S”-shaped TSRs were expressed at lower levels than those with “N”-shape TSRs and those with “B”-shaped TSRs (**Fig. 2b**). In addition, the genes with “S”-shaped TSRs exhibited higher nonsynonymous substitution (*K_a_*)/synonymous substitution (*K_s_*) ratios (i.e., μ values), in comparison with their respective orthologous genes in common bean (*Phaseolus vulgaris*) – a species that diverged from soybean around 17 million years ago (Zhao et al. 2017) – than those with “N”-shaped TSRs and those with “B”-shaped TSRs, suggesting that the former have undergone lower intensities of purifying selection than the latter (**Fig. 2c, Supplemental Table S3, and Supplemental Fig. S3a**).

**Fig. 2.**
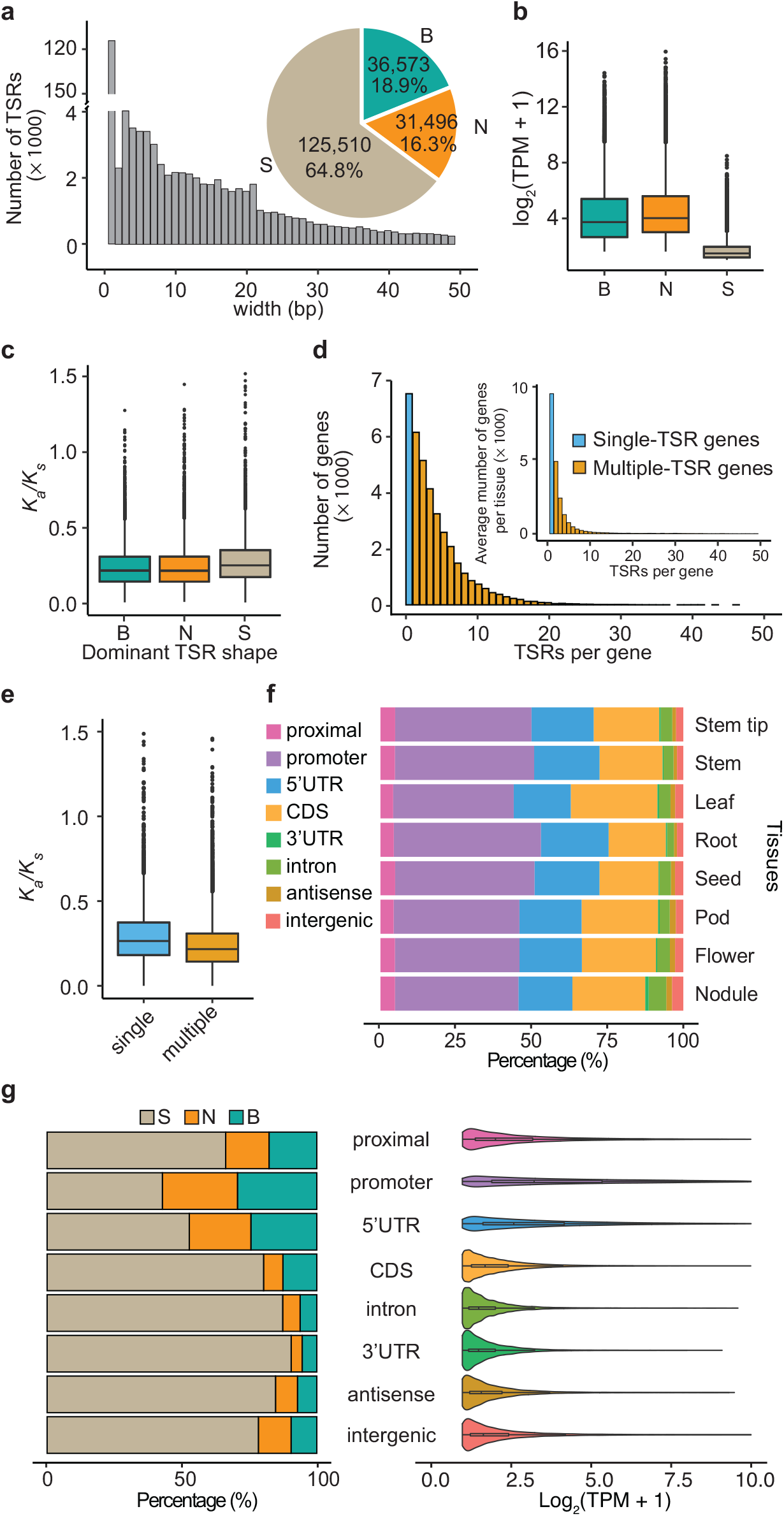
The characteristics of TSRs detected by STRIPE-seq in soybean. **a,** The bar plot displays the length distribution of TSRs. The number and proportion of the three TSR shapes are displayed in the pie chart. Only the length of TSRs that are less than 50 bp were displayed in the bar plot. **b,** The abundance of TSRs in the three shape categories. **c,** The *K_a_/K_s_* value of TSRs in the three shape categories. **d,** The number distribution of TSRs per gene. The outer and inner bar plots display the distribution in total and average of each tissue, respectively. The single and multiple-TSR genes are represented by the blue and orange bars on the plot, respectively. **e,** The *K_a_/K_s_* value of TSRs in single and multiple-TSR genes. “single” and “multiple” represent single-TSR and multiple-TSR genes, respectively. **f,** The relative proportion of TSRs in different annotated features in each tissue. **g,** The relative proportion of the three TSR shapes (left panel) and the abundance of TSRs (right panel) in each feature. “B”, “N”, and “S” represent “B”-, “N”-, and “S”-shaped TSRs, respectively.

Of the 37,911 soybean genes with defined TSR by STRIPE-seq, 30,378 (80.1%) were found to contain multiple TSRs, either within the same tissue or across different tissues, but 33,509 (88.3%) have fewer than ten TSRs (**Fig. 2d**). When TSRs from an individual tissue were analyzed, averagely, 42.2% genes were found to have a single TSR, but 90% of those genes with multiple TSRs had fewer than five TSRs (**Fig. 2d and Supplemental Fig. S4**). Furthermore, genes with multiple TSRs exhibited lower *K_a_*/*K_s_* ratios (**Fig. 2e** and **Supplemental Fig. S3b**), suggesting that they have undergone stronger purifying selection than those with a single TSR.

### Non-canonical TSRs in intragenic regions are widespread and have distinct features from canonical TSRs

In general, TSRs are expected to be located in regulatory regions such as proximal sequences, promoters, or 5’ UTRs, and referred to as canonical TSRs. We found that 97,528 (50.4%) of the TSRs defined in this study fell into the category of canonical TSRs, of which 56.2% were situated in promoter regions annotated in the reference genome (**Fig. 2f**). Interestingly, 84,312 (43.5%) of the TSRs were found in annotated introns, CDSs, or 3’UTRs, dubbed intragenic regions. Strikingly, of these intragenic TSRs, 82.5% were located in CDSs. These CDS-TSRs (C-TSRs) account for 18.9% in roots up to 28.6% in leaves among all TSRs located in regulatory and intragenic regions. (**Fig. 2f**). In addition to the TSRs in regulatory and intragenic regions, 6,907 (∼3.6%) of TSRs were positioned in intergenic regions, likely corresponding to non-coding RNAs or unannotated genes (**Fig. 2f**). We observed that regulatory regions had a greater proportion of “B”-shaped TSRs than intragenic regions, which harbor the highest proportion of “S”-shaped TSRs (**Fig. 2g**). Only in the promoter regions did “S”-shaped TSRs account for <50% of all TSRs (**Fig. 2g**).

We then compared sequences surrounding TSRs in different regions. TATA-box elements were detected at 30 bp upstream of TSRs in the regulatory and antisense, and intergenic regions (**Supplemental Fig. S5**). In contrast, the TATA-box elements were barely seen around intron-TSRs and were not detected around C-TSRs and 3’UTR-TSRs (**Supplemental Fig. S5**). The distribution of the dinucleotide PyPu elements was considerably consistent among different categories of the TSR regions, with the CA and TG predominated. Nevertheless, slight variations were seen, for example, TA was enriched in regulatory regions, whereas GG was enriched in intragenic regions (**Supplemental Fig. S6**). These observations suggest that transcription in CDSs may be initiated via mechanisms distinct from that in canonical promoters located within regulatory regions.

### ATI was shaped by WGD and the subsequent genomic fractionation

Functional divergence of duplicated genes was often reflected by their diverged expression patterns (Zhao et al. 2017). To understand how TSRs of duplicated genes retained after the recent WGD event have evolved to contribute to their functional divergence and how TSRs of duplicated genes and singletons have been shaped by the genomic fractionation process, we analyzed expression patterns and ATI of WGDs and singletons across eight tissues and found the following patterns: *i*) Similar to reported earlier (Zhao et al. 2017), overall the duplicated genes were expressed at higher levels than the singletons (**Fig. 3a and Supplemental Fig. S7**); *ii*) the duplicated genes possessed more TSRs than the singletons (**Fig. 3b, c and Supplemental Fig. S7**); *iii*) the duplicated genes exhibited fewer tissue-specific TSRs than the singletons (**Fig. 3d and Supplemental Fig. S7**). When two members of each of the duplicated gene pairs with detected TSRs were compared, we found that the members expressed at higher levels possessed more TSRs and fewer tissue specific TSRs than their duplicates (**Fig. 3e, f and Supplemental Fig. S7**). These observations suggest that the distribution patterns of TSRs are important indicators of functional conservation and divergence of duplicated genes.

**Fig. 3.**
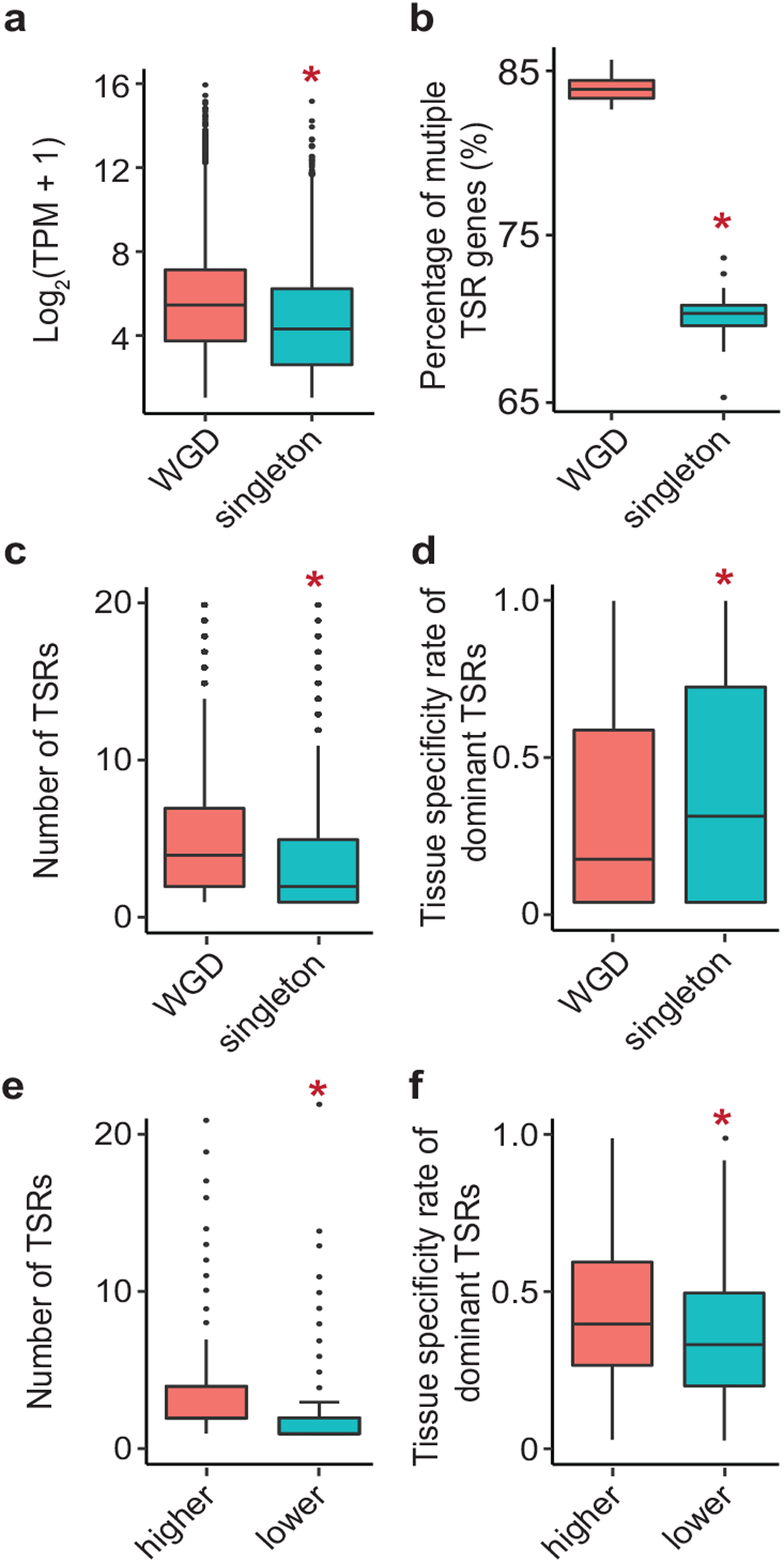
The characteristics of TSR divergence after whole-genome duplication in soybean. **a,** The expression abundance in each gene group. The expression abundances were normalized by TPM and log2 transformed. **b,** The percentage of multiple TSR genes in each gene group. **c,** The number of TSRs in one gene in each gene group. **d,** The tissue-specific rate of dominant TSRs in each gene group. **e,** The number of TSRs in one gene in high-expressed WGD copy and lower-expressed WGD copy. **f,** The tissue-specific rate of dominant TSRs in high-expressed WGD copy and lower-expressed WGD copy. *KS* test was performed to assess statistical significance (\*\**p* value < 0.01).

### Antisense and sense TSRs exhibited similar distribution patterns along genic sequences, and so did bidirectional TSRs

The strand-specific nature of STRIPE-seq allowed us to determine antisense and bidirectional TSRs and their relative abundance. An antisense TSR was defined when a sense TSR existed within the same gene body (**Fig. 4a**). In the eight soybean tissues, we identified 4,574 antisense TSRs, which were assigned to 3,170 genes (**Supplemental Table S4**). These antisense TSRs were predominantly tissue-specific, comprising 85% of all the antisense TSRs. In addition, the antisense TSRs were most frequently found in the CDS regions, harboring 39.5%, on average, across tissues. In contrast, only 5.2% of the antisense TSRs were found in the 5’UTR regions (**Fig. 4b**). Anti-sense TSRs were also observed in intronic sequences, for example, in nodules, we detected an antisense TSR and corresponding transcript in *Glyma.01G009700*, a gene putatively encoding a transmembrane protein, and this antisense TSR originated within the intronic region of the annotated sense transcript (**Fig. 4c**).

**Fig. 4.**
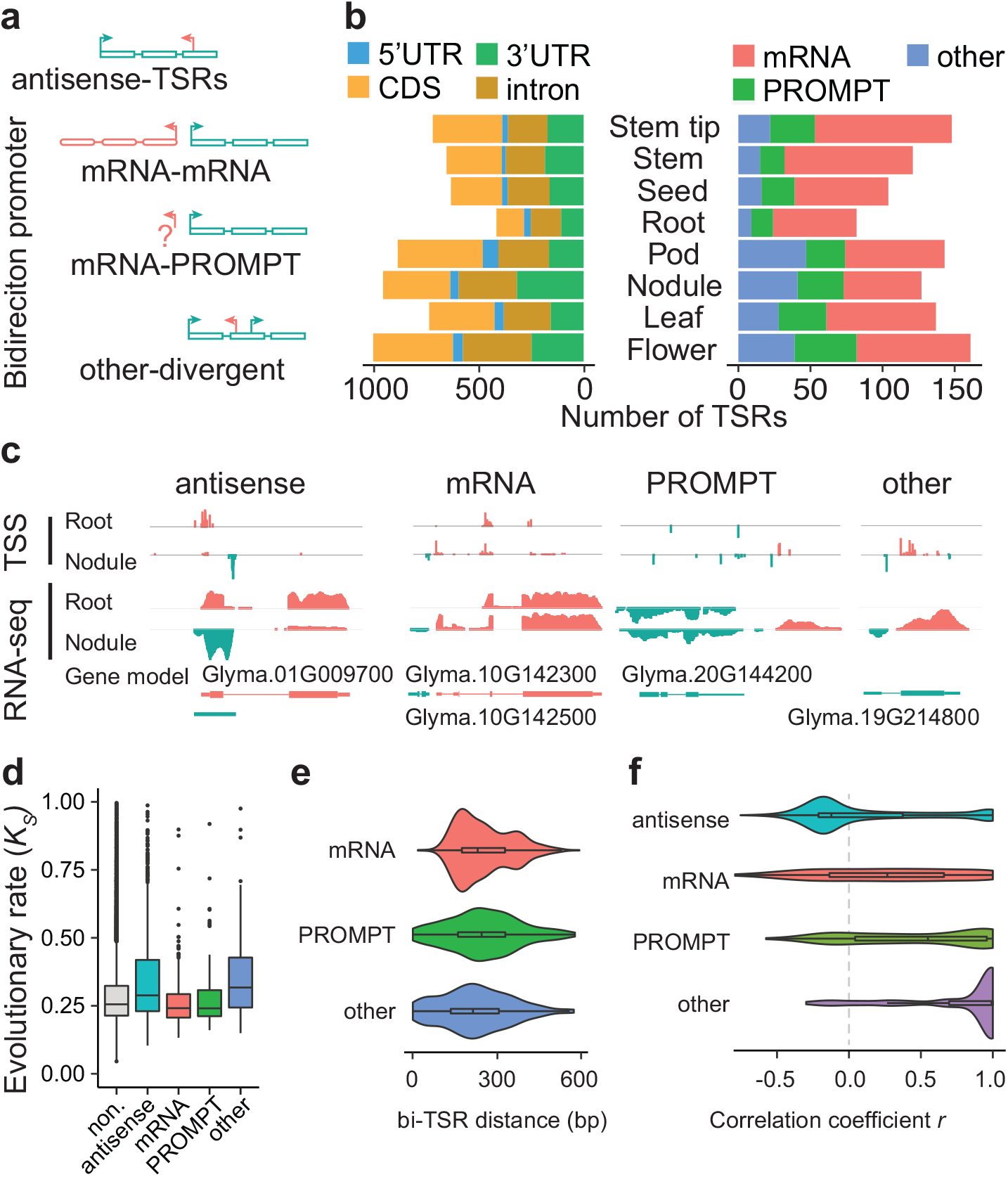
The characteristics of antisense- and bidirectional TSRs in soybean. **a,** Schematic representation of the categories of antisense- and bidirectional TSRs. The rectangle represents the exons of a given gene. The arrows with green and red colors represent the sense- and antisense-TSRs. **b,** The number of antisense-TSRs in annotated gene features and the catteries of bidirectional TSRs in each tissue. **c,** The examples of antisense- and bidirectional TSRs in Root and Nodule. The red and green colors represent the sense and antisense reads relative to the genome. the blue bar at the bottom represents the structure of the gene models. **d,** The evolutionary rate (*Ks*) in genes with different TSR categories. “non.” represents the genes without antisense- and bidirectional TSRs. **e,** The distance distribution of three types of bidirectional TSRs. The x-axis represents the longest distance between two directional TSRs in one bidirectional TSR. **f,** The correlation of antisense-TSR and the sense TSR abundance among tissues in each category. “antisense”, “mRNA”, “PROMPT” and “other” represent bidirectional TSRs belonging to the antisense-TSR, mRNA-mRNA, mRNA-PROMPT, and other-divergent categories, respectively.

Bidirectional TSRs were defined when they were found within 600 bp to initiate transcription in opposite directions (**Fig. 4a**). We identified 506 bidirectional TSRs, including 156 mRNA-mRNA pairs, 162 mRNA-promoter upstream transcripts (PROMPT) pairs, and 184 other-divergence pairs in the eight soybean tissues assigned to 664 genes. (**Fig. 4b and Supplemental Table S5**). These bidirectional TSRs, particularly mRNA-PROMPT and other divergent forms, also showed a high degree of tissue-specificity, (**Fig. 4b and Supplemental Table S5**). We also observed an “mRNA-mRNA” type of TSR interaction between *Glyma.10G142300*, which generates a potential non-coding RNA, and *Glyma.10G142500*, a gene encoding a neurofilament-like protein (**Fig. 4c**). Additionally, an “mRNA-PROMPT” type of TSR was identified in *Glyma.20G144200*, a gene encoding a putative protein belonging to the major facilitator superfamily (**Fig. 4c**). An example of other-diverged type of TSR was found in *Glyma.19G214800*, whose putative ortholog in cotton is involved in fiber expression. The antisense transcript of *Glyma.19G214800* was initiated in its intronic region (**Fig. 4c**). These specific TSRs in nodules were supported by RNA-seq data, which could not precisely define the boundaries of the TSRs though (**Fig. 4c**).

Our analysis also reveals that the genes producing antisense and other divergent TSRs have evolved at an overall faster pace, as reflected by *K_s_*, than the genes without antisense or divergent TSRs (**Fig. 4d and Supplemental Fig. S8a**). Interestingly, the earlier have undergone stronger intensities of purifying selection than the latter (**Supplemental Fig. S8b**), suggesting the functional constraints of these alternative forms of transcripts. On average, the distance between bidirectional TSRs is ∼250 base pairs, close to the length of DNA between two nucleosomes (**Fig. 4e**). Notably, the expression of antisense TSRs was generally negatively correlated with the expression of corresponding sense TSRs, probably minimizing the production of sRNAs from paired transcripts from the same genes. All categories of bidirectional TSR pairs showed positive correlations in abundance of transcripts, suggesting lack of functional interference between the transcripts produced with bidirectional TSRs (**Fig. 4f**).

### ATI in coding sequences tend to be tissue-specific and likely execute tissue-specific functions

To understand functional significance of ATI, we analyzed distribution patterns of all TSRs across the eight tissues. Of the 193,579 TSRs, 62.7% (121,353) were found to be tissue-specific, while 5.5% (10,712) were shared by the eight tissues (**Fig. 5a, and Supplemental Table S2**). Of the 10,712 tissue-specific TSRs, 93.1% were “S”-shaped, while 3.7% were “B”-shaped (**Supplemental Fig. S9**). Interestingly, tissue-specific TSRs were least abundant in roots but most abundant in root nodules - a root specific organs in most legumes (**Supplemental Fig. S10a**), suggesting that the tissue specific TSRs plays an important role in maintaining tissue identity or executing tissue-specific biological functions such as symbiosis. Strikingly, 45.6% of the tissue specific TSRs were found in CDSs and likely enable production of new proteins. By contrast, only 17.7 % of the tissue specific TSRs were in annotated promoter regions (**Fig. 5a**). These tissue-specific C-TSRs and tissue-specific TSRs located in introns and 3’UTRs (together so-called intragenic TSRs) were ∼ four times as many as the tissue specific TSRs located within canonical promoter regions (P-TSRs) (**Fig. 5b and Supplemental Fig. S10b**), illustrating the magnitude of ATI for tissue-specific gene regulation. According to Gene Ontology (GO) analysis of genes with tissue-specific TSRs, several biological processes, such as cell wall organization, growth, secondary metabolism, and transport, were overrepresented (**Supplemental Fig. S11**), highlighting the biological importance of tissue specific TSRs.

**Fig. 5.**
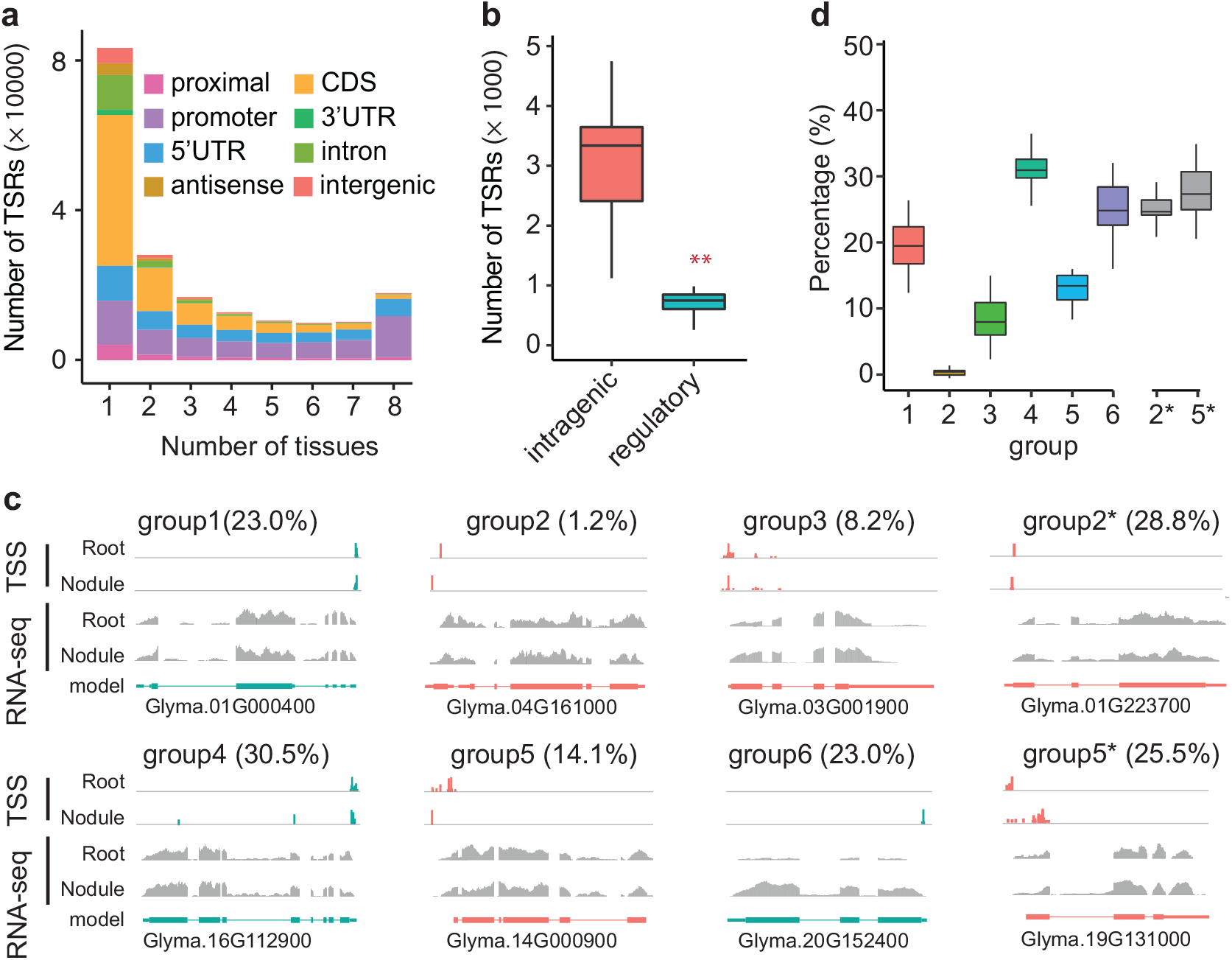
ATI among different tissues in soybean. **a,** The number of TSRs present in a given number of tissues. Different colors represent different annotated features. **b,** The number of tissue-specific TSRs in intragenic and regulatory regions. Different colors represent different annotated features. *KS* test was performed to assess statistical significance (\*\**p* value < 0.01). **c,** Examples of TSRs in root and nodule to illustrate the definition of each group. The red and green colors represent the sense and antisense TSRs relative to the reference genome. the grey bars represent the read coverage of RNAseq in a given gene. the blue bar at the bottom represents the structure of the gene models. The value inside the parentheses represents the percentage of genes in each group in the comparison of root with nodule. **d,** The percentage of genes in each group when comparing between two tissues. The number on the x-axis represents the group number. The group 1-6 percentages are out if the total number of genes while the group2* and 5* percentages are out of the total number of genes in group2 and group5, respectively.

To dive further into the tissue specificity/generality of ATI, we performed pairwise comparisons of TSRs among the eight tissues and then categorized genes into six groups based on the distribution patterns of TSRs between a tissue pair (**Fig. 5c**): Group 1 – genes with a single TSR shared between two tissues compared; Group 2 – genes with a single TSR in each tissue but not shared between two issues compared. Group 3 – genes with multiple TSRs, all shared between the two tissues compared. Group 4 – genes with multiple TSRs, among which the dominant TSR was shared between two tissues compared and at least one TSR was not shared. Group 5 – genes with multiple TSRs, of which the dominant TSRs in each of the two tissues were not shared. Group 6 – genes with TSRs present in only one of the two tissues compared, with no TSR detected in the counterpart. The group 2 genes were least abundant, and the group 4 genes were most abundant. Totaling 15.3% of expressed genes in roots and nodules exhibited ATI between the two tissues, (**Fig. 5c**).

To shed light on potential functional consequences of ATI, we extracted subsets of genes within groups 2 and 5. These two subsets of genes (groups 2* and 5*), accounting for 24.8% and 27.4% of all genes in groups 2 and 5, respectively, were predicted to encoded distinct proteins between two compared tissues due to ATI, respectively. (**Fig. 5c**). These subsets of genes were enriched in processes such as mRNA processing, cell differentiation, and cellular amino acid metabolism (**Fig. 5d, Supplemental Table S6, and Supplemental Fig. S12**). In particular, genes predicted to encode distinct proteins between roots and nodules were enriched in defense response, protein ubiquitination, and transport (**Supplemental Fig. S13**). As exemplified in Figure 5c, *Glyma.01G223700*, a putative USO1-like intercellular protein transport protein gene, was predicted to produce an N-terminus-truncated protein in the roots that is 67-amino acid shorter than the protein produced in the nodules, and the truncation potentially alters its subcellular location. *Glyma.19G131000*, a putative ankyrin repeat family gene, was predicted to generate an N-terminus-truncated protein in the nodules that is 92-amino acid shorter than the protein generated in the roots (**Fig. 5c**), representing functional innovation triggered by ATI in CDSs.

### TSRs in promoters and coding regions exhibited distinct epigenomic features

Genetic and epigenetic features of canonical promoter regions have been well documented. To understand how ATI in intragenic regions is triggered, we examined key histone modifications, (H3K4me3, H3K56ac, H3K36me3, H3K4me1, and H3K27me3) along with histone protein markers (H3), around TSRs with genome-wide chromatin profiles generated from soybean leaves. H3K4me3, H3K56ac, and H3K36me3 are active transcription-associated histone marks, whereas H3K27me3 and H3K4me1 are repressive transcription-associated histone marks (Guenther et al. 2007; Zhao et al. 2019). Overall, the TSRs in promoter regions (P-TSRs) of all expressed genes in the eight tissues were characterized by typical promoter-like chromatin architecture, as represented by overlapped peaks of H3K4me3 and H3K56ac, and H3K36me3 (**Supplemental Fig. S14**). Such a peak was also detected around TSRs identified in the intergenic intervals and around TSRs to produce anti-sense transcripts. By contrast, the intragenic regions exhibited a relatively even distribution of H3K4me3 and H3K56ac, and H3K36me3, which were less abundant in these regions than shown at the peak near the promoter-TSRs (P-TSRs) (**Supplemental Fig. S14**).

Since the C-TSRs identified in the eight tissues were highly tissue-specific and the histone modification profiles were only generated from soybean leaf tissue, we subsequently focused on comparison between the leaf-specific C-TSRs (LsTSRs) and the C-TSRs specific to each of the other seven tissues (OsTSRs) in the context of epigenomic features leaves, with corresponding tissue-specific P-TSRs as controls (**Fig 6a, b, and Supplemental Fig S15**). Similar to the P-TSRs of all expressed genes in the eight tissues, leaf-specific P-TSRs exhibited typical promoter-like chromatin architecture. Interestingly, a relatively low abundance of H3 curve (representing a nucleosome-free region or NFR) was observed at leaf-specific C-TSRs, suggesting that like that in the canonical promoter regions, the chromatin accessibility for transcriptional machinery is critical for the occurrence of transcription initiation at C-TSRs (**Fig. 6c**). Nevertheless, NFRs around C-TSRs were substantially shorter than those around P-TSRs, potentially explaining why fewer TATA-box motifs were found in the former than in the latter (**Fig. 6b and Supplemental Fig. S5**). In addition, a relatively low abundance of H3K4me1, commonly linked to RNA polymerase II (RNAPII) elongation and thus considered as a contributory factor in triggering transcription initiation (Nielsen et al. 2019), was also observed at the C-TSRs (**Fig. 6b**). Furthermore, the C-TSRs were found to be flanked by two peaks of H3K4me3, H3K56ac, and H3K36me3. These observations suggest that the leaf-specific ATI within CDS was epigenetically regulated.

**Fig. 6.**
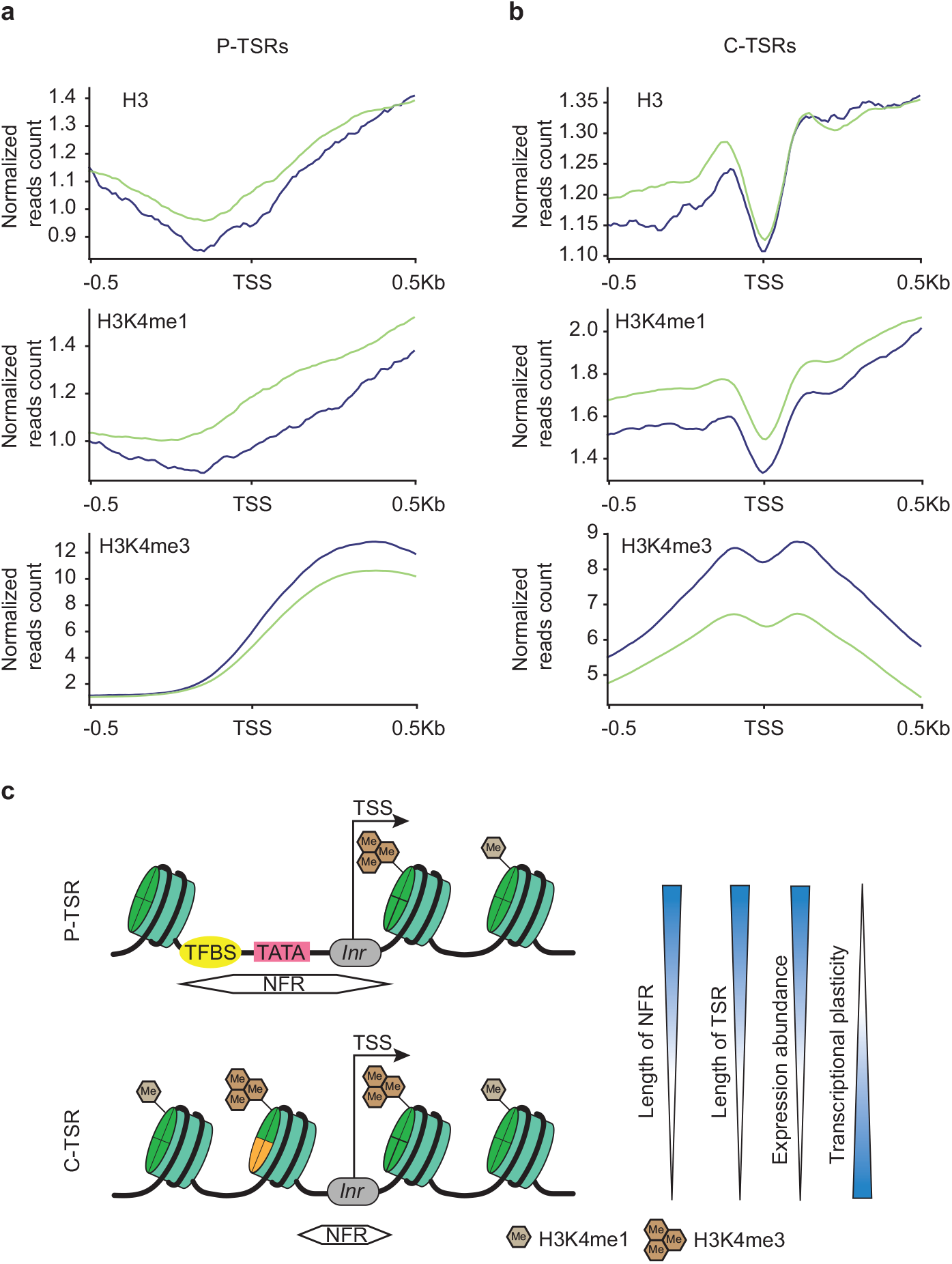
The distributions of histone modifications and variants around leaf-specific and other tissue-specific TSSs located in given feature regions in leaf. **a,** The distribution of markers around P-TSRs. **b,** The distribution of markers around C-TSRs. The TSS was defined as the peak site of the TSR. The lines with different colors represent the different markers. The blue and green lines represent the LsTSRs and OsTSRs, respectively. **c,** Proposed model of nucleosome accessibility and histone modifications around P-TSRs and C-TSRs. The arrow represents the initial location and direction of transcription.

Due to the lack of histone modification profiles from the seven non-leaf tissues, it is unknown if the C-TSRs in each of the seven tissues are also featured by these epigenetic marks. Interestingly, the C-TSRs in each of the seven tissues were found to be located at or adjacent to the H3 curve and H3K4me3, H3K56ac, and H3K36me3 peaks detected in leaves. Further, the active transcription-related markers detected in leaves were found to be more enriched at/around C-TSRs specifically deleted in leaves than C-TSRs in other seven tissues. By contrast, the repressive transcription-related markers detected in leaves were found to be less enriched at/around C-TSRs specifically detected in leaves than C-TSRs in other seven tissues (**Fig. 6a, 6b and Supplemental Figs. S15**). These observations indicate that epigenetic marks are widely preserved across various tissues and exhibit threshold effects on tissue-specific gene expression.

## Discussion

Characterization of TSRs in any plants, including *Arabidopsis thaliana,* remains incomplete and to some extent, inaccurate (Thieffry et al. 2020). Our work demonstrates genomic and epigenomic features shaping alternative transcription initiation by developing a comprehensive plant TSR atlas using the newly developed STRIPE-seq technique and by analyzing their distribution patterns in the context of those well-defined features. Because STRIPE-seq can identify TSSs and simultaneously evaluate relative abundances of transcripts from individual TSSs with as little as 50 ng of total RNA, we were able to effectively identify TSR distribution patterns and assess their tissue specificity at a high degree of precision. In addition to basic findings, our work provides a valuable resource of TSR data that complements existing gene annotation and expression atlases that widely used by the research community, laying a foundation for future studies aimed at unraveling the roles of ATIs in shaping the plasticity of plant phenotypes.

In our analysis of the patterns of transcription initiation, we have observed a blend of conservation and divergence across species. The TSRs situated within the canonical regulatory regions of soybean genes exhibit conserved genomic and epigenomic features including broad TSRs, the presence of TATA-box motifs, relatively long NFRs, and relatively low degrees of tissue specificity in soybean. These characteristics align with those observed in other plant species as well as mammals, underscoring a shared strategy in transcriptional regulation across diverse taxa (Policastro et al. 2020). However, when delving into the expression levels associated with various TSR shapes, soybean presents a divergence. The TSRs with single-nucleotide shapes are linked to lower expression levels in contrast to TSRs with narrow or broader shapes. This finding contradicts to the observation obtained in maize, where genes with single and multiple TSRs exhibit similar expression levels (Mejía-Guerra et al. 2015). A more striking contrast emerges when considering the prevalence of multiple TSRs within a single gene: soybean exhibits a considerable portion of genes with multiple TSRs, a sharp deviation from previous reports in Arabidopsis and cotton, which indicate that the majority of genes harbor a single TSR (Wang et al. 2019; Thieffry et al. 2020). This discrepancy may stem from methodological differences as STRIPE-seq likely offers higher sensitivity than CAGE. Furthermore, the diversity of tissue types and the depth of sequencing employed in our study, encompassing eight distinct tissues, in contrast to the sixteen tissues analyzed in cotton and a single seedling tissue in Arabidopsis, which may influence the observed proportions of TSR diversity (Wang et al. 2019; Thieffry et al. 2020). The proportion of divergent TSRs in soybean is considerably lower, a pattern that does not align with findings in vertebrates, where prevalent bidirectional TSRs were observed and were found to be closely associated with eRNAs (Anderson et al. 2015; Thieffry et al. 2020). Yet, this phenomenon is not exclusive to soybean; it has been observed in both Arabidopsis and maize, suggesting that divergence in TSRs, while less frequent, is likely a common feature in the plant kingdom (Mejía-Guerra et al. 2015; Thieffry et al. 2020). Thieffry et al. (2020) have postulated that the suppression of divergent TSRs in plant genomes may be linked to the unique small RNA landscape. For example, the accumulation of small RNAs around PROMPTs in exosome mutants, especially the 24-nt sRNAs, suggests the involvement of the RNA-directed DNA methylation pathway – a plant-specific DNA methylation pathway absent in mammals (Stroud et al. 2013).

Our study uncovered a vast prevalence of ATI within the soybean genome, accounting for around half of the number of the annotated genes showing evidence of ATI across eight different tissues, and a notable portion of these instances identified in the CDS. This is in stark contrast to maize, where only a small fraction (1.2%) of genes with ATI in the CDS were associated with changes in protein-coding potential (Mejía-Guerra et al. 2015). The implications of ATI are manifold: In UTRs, such variations may result in the gain or loss of regulatory motifs, impacting the regulation of gene expression; Within CDS, they can alter protein function or abundance. For example, ATI has been implicated in significant biological outcomes, such as cancer development in humans (Davuluri et al. 2018). In plants, ATI within CDS regions has been shown to affect protein localization and function. For instance, in maize, two genes encoding a neutral leucine aminopeptidase and a glutathione S-transferase, respectively, exhibit changes in their subcellular localization attributed to ATI (Mejía-Guerra et al. 2015). Predictions for Arabidopsis genes MADR and sAPX indicate similar outcomes (Tokizawa et al. 2017). Furthermore, ATI in CDS can result in protein N-terminus changes, as seen with a maize MYB family gene, which, due to a shifted TSR, loses a zinc-finger domain crucial for its function (Mejía-Guerra et al. 2015). Additionally, the NRT1.2 gene in cotton demonstrated altered nitrate uptake activity owing to ATI, highlighting the functional impact of such changes (Wang et al., 2019). Beyond the CDS, ATI in the UTRs, especially the 5’UTR, can modulate translational efficiency via post-transcriptional mechanisms. or instance, in cotton, ATI has the potential to alter the uORFs of numerous genes (Wang et al. 2019). Taken together, these observations underscore the significance of ATI as a key regulatory mechanism, introducing additional complexity to gene expression patterns in varying tissues, developmental stages, and environmental contexts. Understanding the full spectrum of ATI’s roles remains an emerging and crucial area of research (Batut et al. 2013).

Examination of the TSRs within the soybean genome reveals distinct patterns between those located within CDS and those found in promoter regions. These differences likely originate from the distinct histone occupancy and modification patterns observed in CDS and those in promoter regions, evidenced by substantial variation in nucleosome organization (Soboleva et al. 2014). The more confined NFRs in TSR within CDS, inferred from histone occupancy, imply a distinct regulatory mechanism for transcription initiation. The confined space between nucleosomes surrounding TSRs within CDS accounts for their typically sharper nature compared to those in promoter regions. These subtler NFRs correlate with increased transcriptional plasticity, leading to gene-specific expression across various cell types, developmental stages, and environments (Tirosh and Barkai 2008), explaining the high transcriptional specificity observed in TSRs within CDS. In the promoter regions of transcribed genes, there is an enrichment of H3K4me3, which is closely associated with transcriptional initiation (Guenther et al. 2007). On the other hand, H3K4me1, a marker of RNA polymerase II elongation, is underrepresented in TSRs within CDS in our study. Nielsen et al. reported that H3K4me1 is necessary to repress intragenic initiation of RNA polymerase II transcription in Arabidopsis (Nielsen et al. 2019). The transcription initiation at TSRs within CDS is downregulated by reduced promoter-specific epigenetic markings and increased markings associated with RNA polymerase II elongation (Nielsen et al. 2019). This could explain why highly expressed TSRs within CDSs are associated with lower H3K4me1 signals but higher H3K4me3 signals in soybean. Notably, our observations show consistent chromatin architecture patterns in leaves at specific TSRs in the leaf versus those in other tissues, though with intensified signals for active markers and diminished signals for repressive markers. This implies that TSR chromatin architecture functions as a persistent ‘door’, poised to unlock and fine-tune transcriptional initiation among tissues, with specific epigenetic markers acting as the ‘key’. However, we cannot discount the potential influence of other histone modification markers, like H3K27me3, H3K56ac, and H3K36me3, on transcriptional initiation. Further investigation is needed to fully comprehend their roles in this intricate regulatory landscape.

This study explores alternative transcription initiation (ATI) in soybean, revealing its significant role in regulating gene expression across various tissues, particularly within coding sequences. This extensive influence of ATI underscores its importance in altering protein functionality and prevalence, contributing to soybean’s adaptability and phenotypic diversity. A key finding is the distinct pattern of TSRs in coding regions compared to regulatory regions, characterized by variations in numbers and abundance of TSSs, presence/absence of TATA-boxes, lengths of NFRs, epigenetic features, suggesting a complex interplay between genetic and epigenetic factors, as a finely tuned mechanism for gene expression. This study not only enhances our understanding of transcriptional complexity in an economically important crop but also provides a valuable set of TSRs for further investigation of biological significance of ATI.

## Methods

### Plant Growth Conditions and Material Collection

Soybean (*Glycine max*) cultivar Williams 82 was grown in the greenhouse under 16 h light/8 h dark period. Eight tissues, including root, nodule, stem, stem tip, leaf, flower, pod, and seed were collected at the development stages of trefoil, flowering, and seed development. *Bradyrhizobium diazoefficiens* USDA 110 was used for soybean root inoculation, the nodules on inoculated roots 20 days post inoculation (dpi) and uninoculated roots at the same stage were prepared and harvested as previously described (Ren et al. 2019). Each sample was collected from at least eight individual plants, and the mixed bundle was quickly frozen in liquid nitrogen for RNA isolation.

### RNA isolation, DNA & rRNA depletion

The total RNA was isolated using the TRIzol reagent (Invitrogen/Life Technologies, CA) according to the manufacturer’s instructions. To remove the residual DNA, the extracted RNA was treated with TURBO DNA-free Kit (Invitrogen). The nodule-derived RNA was treated with NEBNext rRNA Depletion Kit (Bacteria) (NEW ENGLAND BioLabs) and RiboMinus Plant Kit for RNA-Seq (Invitrogen) sequentially, while the RNA from seven other tissues was treated with RiboMinus Plant Kit for RNA-Seq (Invitrogen) to deplete the rRNAs.

### STRIPE-seq library construction and sequencing

STRIPE libraries were constructed as previously described and a step-by-step protocol can be found at protocols.io (https://www.protocols.io/view/stripe-seq-library-construction-bdtri6m6) (Policastro et al. 2020). Briefly, to reduce the proportion of uncapped RNA, the DNA & rRNA depleted RNA samples were treated with Terminator 5’-Phosphate-Dependent Exonuclease (Lucigen). After a one-hour incubation, template-switching reverse transcription (TSRT) was performed using a unique barcoded reverse transcription oligo (RTO) per sample, followed by the addition of a unique molecular identifier (UMI)-containing, 5’-biotin-modified template-switching oligo (TSO) with three 3’ riboguanosines. Library PCR is then performed using the TSRT product as input, which ensures that TruSeq adapters are present on both sides of the insert. According to the initial amount of the DNA & rRNA depleted RNA samples, 16-20 cycles of PCR with the forward library oligo (FLO) and reverse library oligo (RLO) were performed in this step. Solid phase reversible immobilization (SPRI) bead size selection is used to remove fragments that are outside the ideal size. Final libraries distributed between 250 to 750 bp with a total amount of 25 to 100 ng were sequenced using the Illumina Novaseq 6000 platform to generate the 150 bp paired-end reads at UC Davis Sequencing Center (Novogene Corporation Inc.). See **Supplemental Table S7** for RTO, TSO, FLO and RLO sequences.

### Processing of STRIPE-seq data

First the raw read quality was estimated with the “fastQC” program (version 0.11.9, https://www.bioinformatics.babraham.ac.uk/projects/fastqc/). The paired 1 read set was used for the following analysis because the strand of reads is the sense direction of RNA. The adapter sequences and poly “G” sequences were trimmed with the “cutadapt” program (version 2.5) (Martin 2021). trimmed reads were removed if they were less than 50 nucleotides or lacked a TATAGGG sequence pattern. Then, the PCR redundant reads were removed by taking advantage of UMI sequences using the “fasx_collapser” function in the “fastx-toolkit” program (version 0.0.14, http://hannonlab.cshl.edu/fastx_toolkit/index.html). The TATAGGG structure of selected reads was trimmed by using the “cutadapt” program. To remove potential rRNA contamination in the data, we collected all the rRNA sequences from the soybean reference genome (version2.1, phytozome) and NCBI database (nt., ftp.ncbi.nlm.nih.gov/blast/db/) (Schmutz et al. 2010). The cleaned reads were mapped to the rRNA sequences using “hisat2” with default parameters. We recalled the reads which unmap to rRNAs. We further mapped all the recalled reads to the soybean reference genome using “hisat2” and kept the records with mapping quality larger than 30 using the “SAMtools” program (version 1.8) (Li et al. 2009). For nodule, which is a symbiotic organ, we additionally removed the sequence from the *Bradyrhizobium diazoefficiens* USDA 110 genome using the above programs (Kaneko et al. 2002).

### Identification, quantification, and annotation of TSRs

First, we converted the bam files into strand-specific bigwig files which were used as input files for identification of transcription start sites (TSSs) using the “CAGEfightR” package (version 1.12.0) in Bioconductor (https://bioconductor.org/) (Thodberg et al. 2019; Gentleman et al. 2004). We only kept the supported TSSs with at least one TPM after combining the abundance of all tissues. Adjacent TSSs were clustered and merged to TSRs if the distance dropped to 20 bp or less. The abundance of TSRs was quantified by the sum of TPM values of all support TSSs in one tissue. the peak location was defined as the position of TSS with maximum abundance in TSRs. To annotate the TSRs in the soybean genome, the Williams 82 reference annotation file (gff3, Phytozome) was used to constructing a database for the following analysis by using the “makeTxDbFromGFF” function in the “GenomicFeatures” (version 1.44.2) package in Bioconductor (Goodstein et al. 2012; Lawrence et al. 2013). The promoter region was defined as the region from 100 bp upstream to 100bp downstream of the annotated TSS in the annotation file. The proximal regions were defined as the 400bp upstream of the promoter regions. The antisense TSRs were annotated if the detected TSR was located in the gene body region with reversed direction relative to the annotated transcriptional direction. The overlapping regions were annotated based on the priority annotation level as illustrated in **Figure 1a**. The location of TSRs was defined as intergenic if no annotated features overlapped with them. TSRs were defined as bidirectional if the two reverse directions of the TSRs were located within the 600 bp region. All TSSs can be accessed through our genome browser portal. They can be visited and explored at http://xtlab.hzau.edu.cn/jbrowser/ (**Supplemental Fig. S16**). The portal is built using the Jbrowse2 framework (Diesh et al. 2023).

### Estimation of evolutionary distance and selection constraint

The syntenic homologous gene pairs between soybean (*Glycine max*) and common bean (*Phaseolus Vulgaris*) were defined in previously published data sets (Zhao et al. 2017). To calculate accurate *K_a_*, *K_s_*, and *K_a_/K_s_*, we first performed coding sequence alignment with the “MUSCLE” program (version 3.8.31) using the default parameters (Edgar 2004). the “yn00” and “baseml” modules in the “PAML” program (version 4.8) were processed to calculate all three evolutionary parameters (Yang 2007).

### Sequencing and processing of RNA-seq

Total RNAs of nodule and root were processed using the Epicentre Ribo-zero™ rRNA Removal Kit (Epicentre, USA) to deplete ribosomal RNAs, and the processed RNA samples were used to construct RNA-seq libraries using the NEBNext® Ultra™ Directional RNA Library Prep Kit for Illumina® (NEB, USA). Then the RNA libraries were sequenced using the Illumina HiSeq 4000 platform to generate 150bp paired-end reads. The raw reads were processed with the “fastx-toolkit” program for the removal of low-quality reads and adaptor sequences, and the processed reads from each library were subsequently mapped to the soybean reference genomes using “Hisat2” (version 2.1.0) with the default parameters (Kim et al. 2019). The reads with map quality > 30 were extracted for the following analysis using “SAMtools” (version 1.8). The BAM files were visualized using the “IGV” software (version 2.8.13) (Thorvaldsdottir et al. 2013).

### Processing of ChIP-seq

The ChIP-seq of histone modifications of leaf tissue in soybean was collected from the previous study (**Supplemental Table S8**), and included H3, H3K4me1, H3K4me3, H3K27me3, H2A.Z, H3K56ac, and H3K36me3 (Lu et al. 2019). The reads were pre-processed with the “fastx-toolkit” program. The clean reads were mapped to the soybean reference genome using the “BWA” program with default parameters. We only kept the reads with map quality above 30 using the “SAMtools” program. Then, the BAM files were converted to BigWig files using “BamCoverage” with the parameters (--ignoreDuplicates --normalizeUsing RPGC --effectiveGenomeSize 990741540). The distribution of histone modification signals surrounding the given TSSs was visualized using the “computerMatrix” and “plotProfiles” programs.

### Gene ontology enrichment analysis

Gene ontology enrichment analysis of a given gene group, such as genes with tissue-specific TSRs, was performed and visualized with the “clusterProfiler” package (version 4.0.5) in Bioconductor (Gentleman et al. 2004; Yu et al. 2012). The genes with detected TSRs were selected as the background gene list for the enrichment analysis. GO term information for soybean was extracted from the gene annotation files in the Phytozome database (V12). GO terms with FDR-adjusted *p*-values < 0.05 were considered to be significantly enriched.

## Data Access

The raw read sequences of STRIPE-seq, RNA-seq, and small RNA-seq generated by this study are deposited in the National Center for Biotechnology Information Sequence Read Achieve (http://www.ncbi.nlm.nih.gov/sra/) under the accession number PRJNA757465 and PRJNA757638. Details of tissues, and reprocessed public datasets, include methylC-seq and ChIP-seq for histone modifications, were listed in Supplemental Table S7.

## Acknowledgements

This work was partially supported by the Agriculture and Food Research Initiative of the USDA National Institute of Food and Agriculture (grants 2021-67013-33722, and 2022-67013-37037), National Science Foundation Plant Biotic Interaction (grant 2128023), and the Purdue AgSEED program to JM.

## Author contributions

JM, XW and JD designed the research; XW, CBC and WF analyzed the data; JD performed the research; JM and XW wrote the manuscript.

